# YAP regulates periosteal expansion in fracture repair

**DOI:** 10.1101/2024.12.23.630086

**Authors:** Madhura P Nijsure, Brendan Tobin, Dakota L Jones, Annemarie Lang, Grey Hallström, Miriam Baitner, Gabrielle I Tanner, Yasaman Moharrer, Christopher J Panebianco, Elizabeth G Seidl, Nathaniel A Dyment, Gregory L Szeto, Levi Wood, Joel D Boerckel

## Abstract

Bone fracture repair initiates by periosteal expansion. The periosteum is typically quiescent, but upon fracture, periosteal cells proliferate and contribute to bone fracture repair. The expansion of the periosteum is regulated by gene transcription; however, the molecular mechanisms behind periosteal expansion are unclear. Here, we show that Yes-Associated Protein (YAP) and transcriptional co-activator with PDZ-binding motif (TAZ) mediate periosteal expansion and periosteal cell proliferation. Bone fracture increases the number of YAP-expressing periosteal cells, and deletion of YAP and TAZ from Osterix (Osx) expressing cells impairs early periosteal expansion. Mechanistically, YAP regulates both ‘cell-intrinsic’ and ‘cell-extrinsic’ factors that allow for periosteal expansion. Specifically, we identified Bone Morphogenetic Protein 4 (BMP4) as a cell extrinsic factor regulated by YAP, that rescues the impairment of periosteal expansion upon YAP/TAZ deletion. Together, these data establish YAP mediated transcriptional mechanisms that induce periosteal expansion in the early stages of fracture repair and provide new putative targets for therapeutic interventions.

## Introduction

Bone fractures heal because the skeleton holds a pool of progenitor cells that reside in the periosteum. While we have known for a long time that the periosteum is critical for proper healing outcomes, the underlying mechanisms are unclear. Periosteal cells activate upon injury, proliferating and expanding the periosteum to initiate fracture repair^1^. These cellular functions are coordinated by transcriptional programs. The goal of this study is to understand the transcriptional mechanisms that enable quiescent periosteal cells to activate and expand.

The periosteum contains a diversity of cells, whose function may be indicated by the transcription factors they express. Osterix (Osx), encoded by *Sp7*, is a transcription factor essential in bone development^2^ and repair^3^, and marks a subset of periosteal cells. Initiation of fracture repair specifically requires the expansion of Osx^+^ progenitor cells, whose activity is determined by transcriptional regulators that integrate environmental cues. Among these regulators are Yes Associated Protein (YAP) and its paralog transcriptional coactivator with PDZ binding motif (TAZ). We previously identified YAP and TAZ as important mediators of periosteal expansion in Osx^+^ cells. Conditional deletion of YAP and TAZ from Osx^+^ cells impairs periosteal expansion in the early stages of fracture repair^4^. YAP and TAZ lack DNA-binding domains and rely on transcription factors like Transcriptional enhanced associated domain (TEADs)^5^, which can bind to specific DNA motifs, to initiate transcription. Together, YAP/TAZ and TEAD regulate genes such as *Ctgf* and *Cyr61*. The specific molecular mechanisms through which YAP and TAZ regulate periosteal expansion are unknown. We aim to identify YAP/TAZ-associated DNA binding transcription factors, as well as their transcriptional targets in periosteal cells.

Here, we demonstrate that YAP and TAZ regulate periosteal expansion during the early stages of fracture repair. Using a YAP gain-of-function model, we identified immediate transcriptional targets and transcription factor binding partners. Among these targets, we identified Bone Morphogenetic Protein 4 (*Bmp4*) as a YAP-TEAD target gene. Exogenous BMP4 delivery improved periosteal expansion in YAP/TAZ knockout mice. Together, these data establish YAP-mediated transcriptional programs that promote periosteal expansion.

## Results

### YAP/TAZ deletion impairs periosteal expansion in the early stages of fracture repair

Bone fracture repair initiates by periosteal cell activation, resulting in periosteal expansion at 4 days post-fracture (DPF) (**Fig. 1 A**). To determine the roles of YAP and TAZ in the fracture-activated periosteum, we inducibly deleted YAP and TAZ from Osterix/Sp7 (Osx)-expressing cells, using Osx1-GFP::Cre; YAP^fl/fl^TAZ^fl/fl^ (YAP/TAZcKO^Osx^)^6^ (**Fig. 1 B**). First, we probed for YAP protein, DAPI and EdU to mark proliferating cells in the periosteum (**Fig. 1 C**). In the wildtype (YAP^fl/fl^TAZ^fl/fl^ or WT^fl/fl^) periosteum, fracture increased the number and frequency of nuclear YAP-positive cells at 4 days post-fracture (DPF) (**Fig. 1 D, E**), suggesting that fracture activates YAP/TAZ signaling in the periosteum. Gray bar indicates levels in uninjured WT^fl/fl^ mice. Osx-conditional YAP/TAZ deletion reduced the number and frequency of YAP^+^ cells. In wild type mice, fracture caused periosteal expansion, from 10-20 μm in the intact periosteum to 30-80 μm at 4 DPF. Osx-conditional YAP/TAZ deletion impaired periosteal expansion (**Fig. 1 F**) and decreased total periosteal cellularity (**Fig. 1 G**).

**Figure 1:**
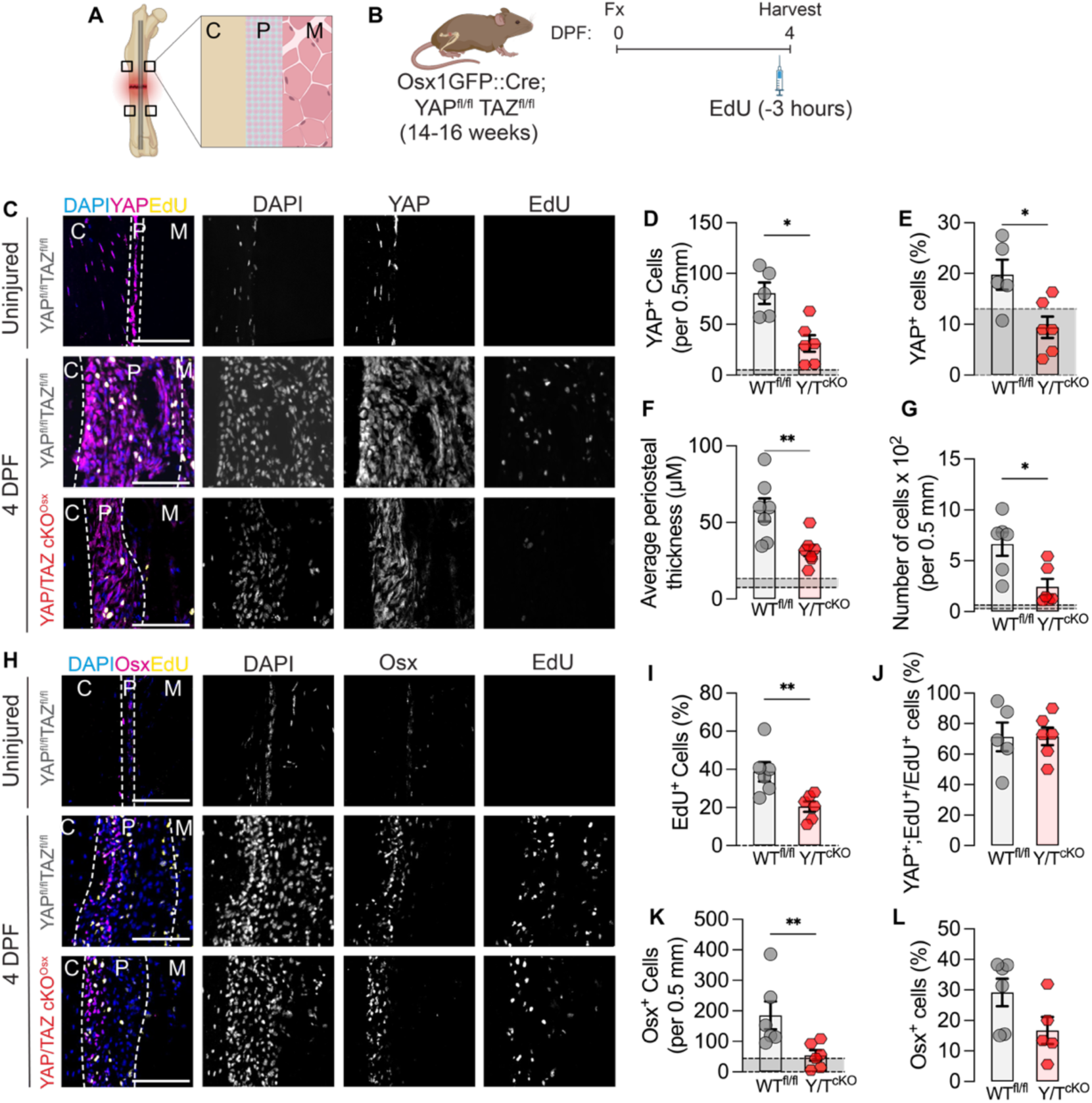
YAP/TAZ deletion from Osx expressing cells impairs periosteal expansion. **A.** Schematic for evaluation of periosteal expansion at 4 Days Post Fracture (DPF). **B.** Timeline for mouse femoral osteotomy, followed by EdU injections at 4 DPF, 3 hours prior to euthanasia. **C.** Staining of nuclei (DAPI), proliferating cells (EdU) and YAP protein. Quantification of **D.** YAP^+^ cells per length of the periosteum, **E.** Percentage of YAP^+^ cells in the periosteum, **F**. Average periosteal thickness, **G.** Number of cells per length of the periosteum after Osx-conditional YAP/TAZ deletion. **H.** Staining of nuclei (DAPI), proliferating cells (EdU) and Osterix (Osx) protein. Quantification of **I**. Percentage of EdU^+^ cells, **J.** Percentage of proliferative YAP^+^ cells, **K.** Osx^+^ cells per length of the periosteum and **L.** Percentage of Osx^+^ cells in the periosteum after Osx-conditional YAP/TAZ deletion. All scale bars are 100 μM. C. Cortical Bone, P = Periosteum, M = Muscle. Gray bars indicate levels in uninjured bones in WT mice. ‘*’: p<0.05, ‘**’: p<0.01.

To quantify periosteal cell proliferation, we administered EdU 3 hours before euthanasia (**Fig. 1 B**). Next, we probed for Osx protein in the periosteum (**Fig. 1 H**). Osx-conditional YAP/TAZ deletion significantly reduced periosteal proliferation (**Fig. 1 I**). Of the EdU^+^ proliferating cells, approximately 80% exhibited nuclear YAP, in both wild type and YAP/TAZ cKO^Osx^ groups, suggesting that the reduction in proliferating cells corresponded to the reduction in YAP^+^ cells (**Fig. 1 J**). YAP/TAZ deletion also significantly reduced the number of Osx^+^ cells (**Fig. 1 K**), while differences in the percentage of Osx^+^ cells were not significant (**Fig. 1 L**). Together, these data indicate that fracture activates YAP in the expanding periosteum and that YAP/TAZ deletion from Osx-expressing cells impaired the expansion of periosteal cells.

### Temporal dynamics of YAP target gene induction

Next, to evaluate transcriptional targets of fracture-activated YAP in periosteal cells, we used a gain-of-function approach. We used a B6.C-tg(CMV-Cre)1Cgn/J;tetO-YAP^S127A^;Gt(ROSA)26^Sortm1(rtTA,EGFP)Nagy/J^ mouse model^7–9^ which, upon doxycycline administration, expresses a mutated, constitutively active form of human YAP (YAP^S127A^, or YAP^CA^). We isolated periosteal cells^10^ at 4 DPF and treated *in vitro* with 1μM Doxycycline to induce YAP^CA^ expression (**Fig. 2 A**). 24 hours of Doxycycline treatment significantly induced *Yap* mRNA expression and upregulated canonical YAP-TEAD target genes, *Ctgf* and *Cyr61*^5^. Similarly, at 24 hours, Doxycycline treatment significantly increased YAP protein levels, without altering TAZ protein (**Fig. 2 E, F**, **Suppl. Fig. 1 A**). 24 hours of YAP^CA^ expression also significantly increased cell proliferation *in vitro* (**Fig. 2 G**).

**Figure 2:**
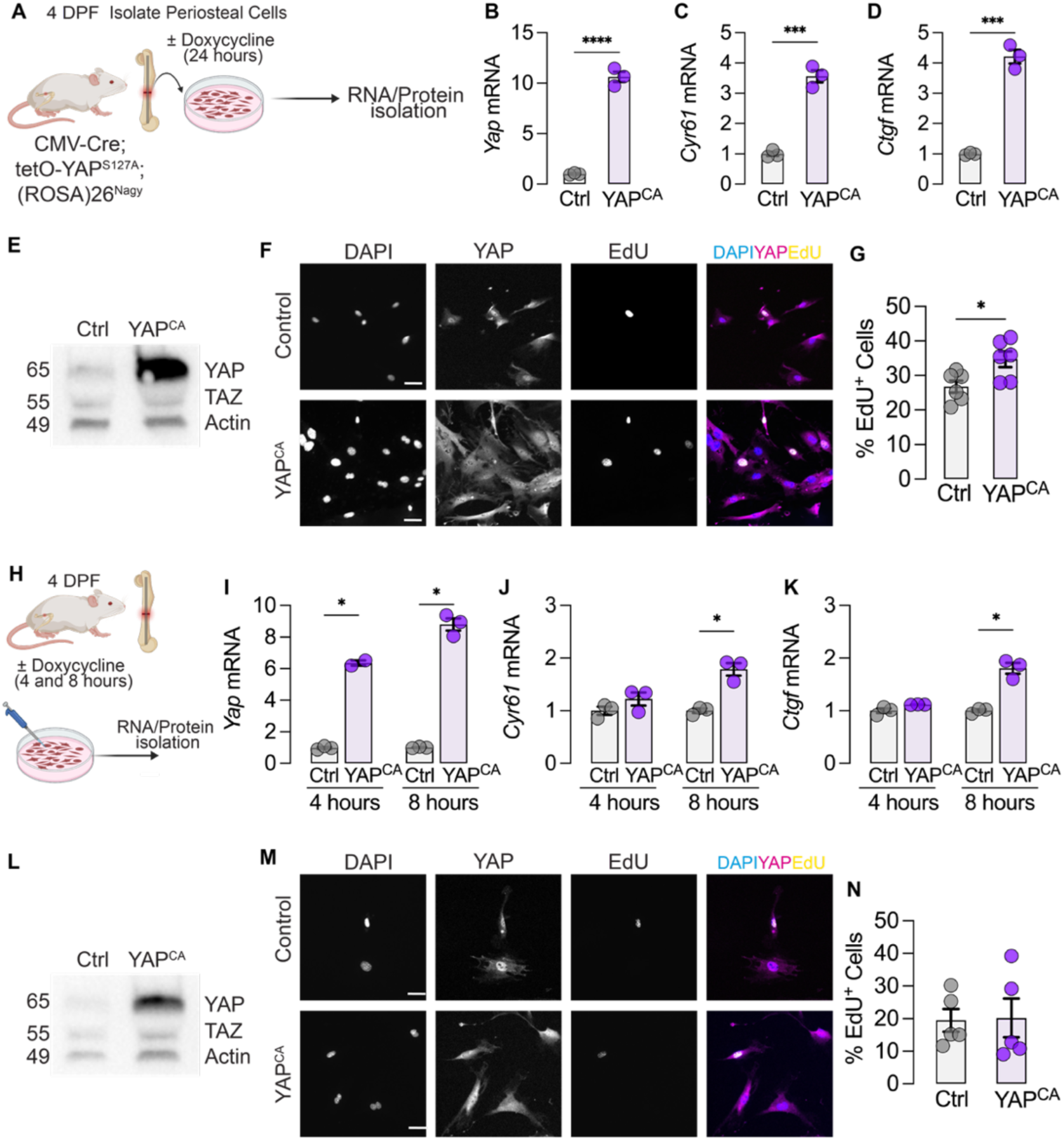
Establishment of a gain-of-function model to identify early YAP target gene. **A.** Schematic for femoral osteotomies and periosteal cell isolation followed by doxycycline treatment *in vitro*. Quantification of mRNA for *YAP* (**B**), *Cyr61* (**C**) and *Ctgf* (**D**). Western blot for quantification of YAP, TAZ and Actin protein levels. Molecular weights are in kDa. **F**. Immunofluorescence staining for DAPI, YAP and EdU in untreated cells and YAP^CA^ expressing cells. **G**. Quantification of the fraction of EdU^+^ cells after 24 hours of YAP^CA^ expression. **H**. Schematic for periosteal cell isolation and RNA/protein isolation after 4 and 8 hours of YAP^CA^ expression. Quantification of mRNA of *YAP* (**I**), *Cyr61* (**J**), *CTGF* (**K**). **N.** Western blot quantification of YAP/TAZ and Actin protein levels. Mol wt are in kDa. **L.** Immunofluorescence staining for DAPI, EdU and YAP 8 hours after YAP^CA^ expression. **M.** Quantification of the fraction of EdU^+^ cells after 8 hours of YAP^CA^ expression. Scale bars = 10 μM. ‘*’:p<0.05, ‘***’: p<0.001, ‘****’:p<0.0001.

YAP activation immediately induces direct target gene expression, but long-term activation can lead to secondary changes in transcription^11,12^. Therefore, to identify early transcriptional targets, without the interference of secondary changes, we evaluated *Yap* expression and YAP-target gene expression at 4 and 8 hours after Doxycycline treatment (**Fig. 2 H**). At 4 hours, Doxycycline treatment significantly induced *Yap* mRNA expression (**Fig. 2 I**), but did not yet induce YAP-target gene expression (**Fig. 2 J, K**). However, by 8 hours, Doxycycline treatment significantly induced both *Yap* and YAP target genes, *Cyr61* and *Ctgf* (**Fig. 2 L, M**) and significantly increased YAP protein levels (**Fig. 2 L, M**, **Suppl. Fig. 1 A**). Additionally, 8 hours of YAP^CA^ expression did not yet induce cell proliferation (**Fig. 2 N**). Based on these findings, we selected 8 hours of Doxycycline treatment as our timepoint to evaluate the YAP-induced transcriptional landscape.

### YAP regulates cell intrinsic and extrinsic pathways that drive periosteal expansion

To this end, we performed mRNA-sequencing and ATAC-sequencing of periosteal cells after 8 hours of YAP^CA^ expression. Bulk mRNA-sequencing of YAP^CA^ cells, compared to untreated controls, identified 376 upregulated genes, and 275 downregulated genes (**Fig. 3 A**). Doxycycline treatment did not significantly alter YAP-target gene expression in periosteal cells isolated from C57BL/6J mice (**Suppl. Fig. 1 B**).

**Figure 3:**
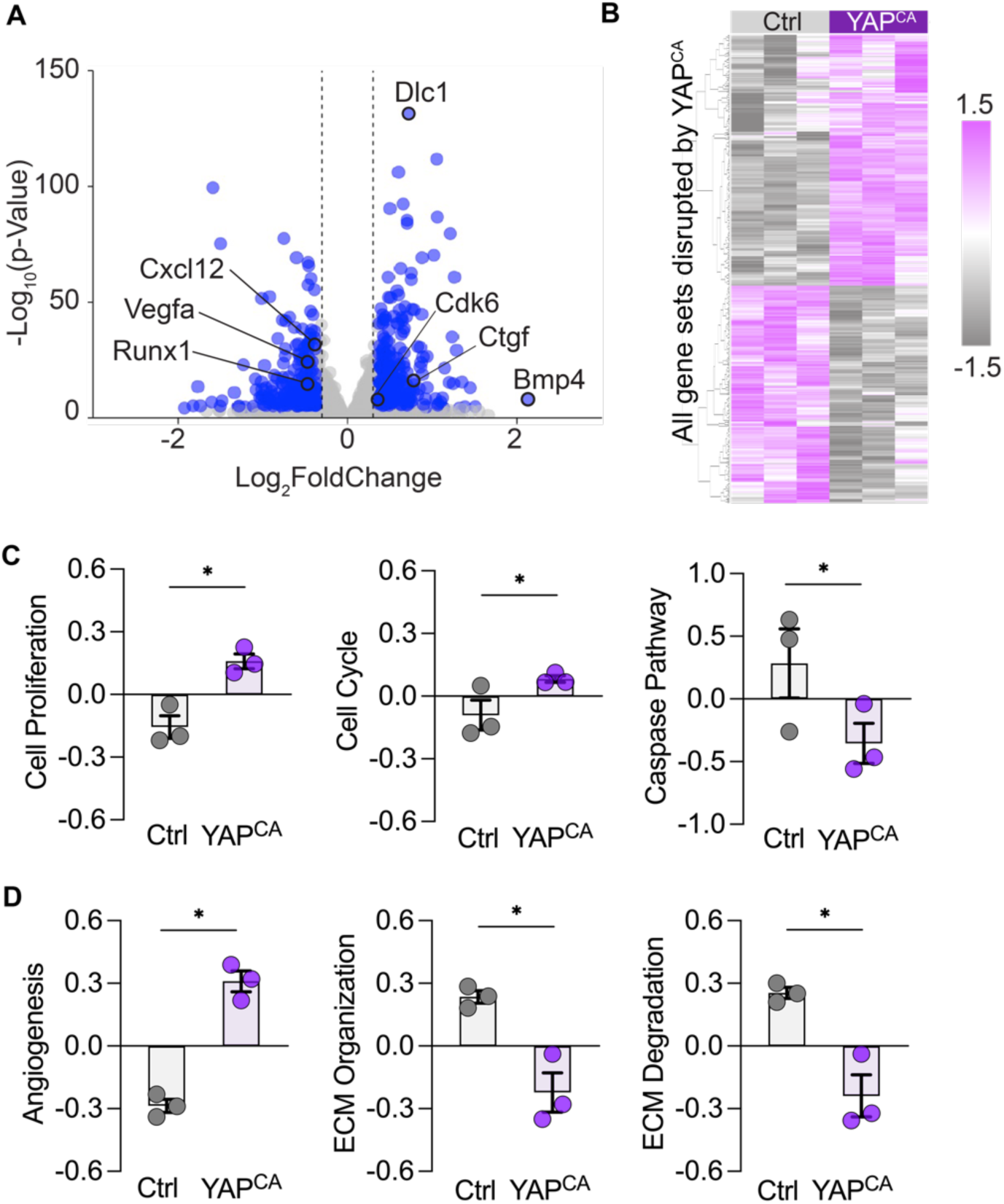
YAP regulates cell intrinsic and extrinsic pathways that drive periosteal expansion. **A.** Volcano plot indicating important genes up- or down-regulated after 8 hours of YAP^CA^ expression. Blue dots indicate genes with −0.3>Log2FoldChange>0.3 and p value less than 0.05. **B.** Heat map indicating enrichment scores for all pathways dysregulated by YAP^CA^. **C.** 261 selected gene sets that could play a role in periosteal expansion. Individual enrichment scores of **D.** cell intrinsic pathways and **E.** cell extrinsic pathways. ‘*’ padj<0.05.

Quantitative gene set variation analysis^13^ revealed 553 significantly differentially regulated gene sets after YAP activation (**Fig. 3 B**). We selected 261 gene sets most relevant to periosteal expansion during fracture repair (**Fig. 3 C**). YAP activation upregulated “cell-intrinsic” pathways involved in proliferation, cell cycle regulation, self-renewal, and protection from apoptosis (**Fig. 3 D**), and “cell-extrinsic” pathways, which may influence neighboring cells and tissues, such as angiogenesis and ECM organization (**Fig. 3 E**). With respect to target genes, YAP activation regulated mRNAs that encode both “intrinsic factors” (such as *Cdk6*), and “extrinsic factors” such as *Bmp4* and *Cxcl12*. Together, these data suggest that YAP activation regulated genes and gene sets for both cell-intrinsic and -extrinsic pathways that could drive periosteal expansion and proliferation.

### YAP/TAZ deletion *in vivo* alters both cell-intrinsic and -extrinsic functions in periosteal cells

Therefore, to test whether YAP/TAZ signaling regulates both cell-intrinsic and -extrinsic activity during fracture repair *in vivo*, we next evaluated the effects of Osx-conditional YAP/TAZ deletion on both Osx^+^ and Osx^−^ cells in the periosteum. We hypothesized that if there is indeed a change in cell-extrinsic factors due to YAP/TAZ deletion, we would see this effect on the Osx^−^ cell population.

Consistently, Osx-Conditional YAP/TAZ deletion decreased both Osx^+^ and Osx^−^ cells (**Suppl. Fig. 1 D**). We observed a gradient of Osterix-expressing cells in the periosteum, with Osx^+^ cells located closest to the bone and Osx^−^ cells positioned farther away. Therefore, we evaluated the spatial distribution of Osx^+^ and Osx^−^ cells in the periosteum (**Fig. 4 A**). This is consistent with a recent report that demarcates the ‘inner’ and ‘outer’ layers of the periosteum as Osx^+^ and ‘Osx^−^ respectively^14^. Endogenous Osx colocalized with Cre::GFP expression, confirming Cre expression to be exclusively restricted to the inner layer (**Fig. 4 B**, **Suppl. Fig. 1 F**).

**Figure 4:**
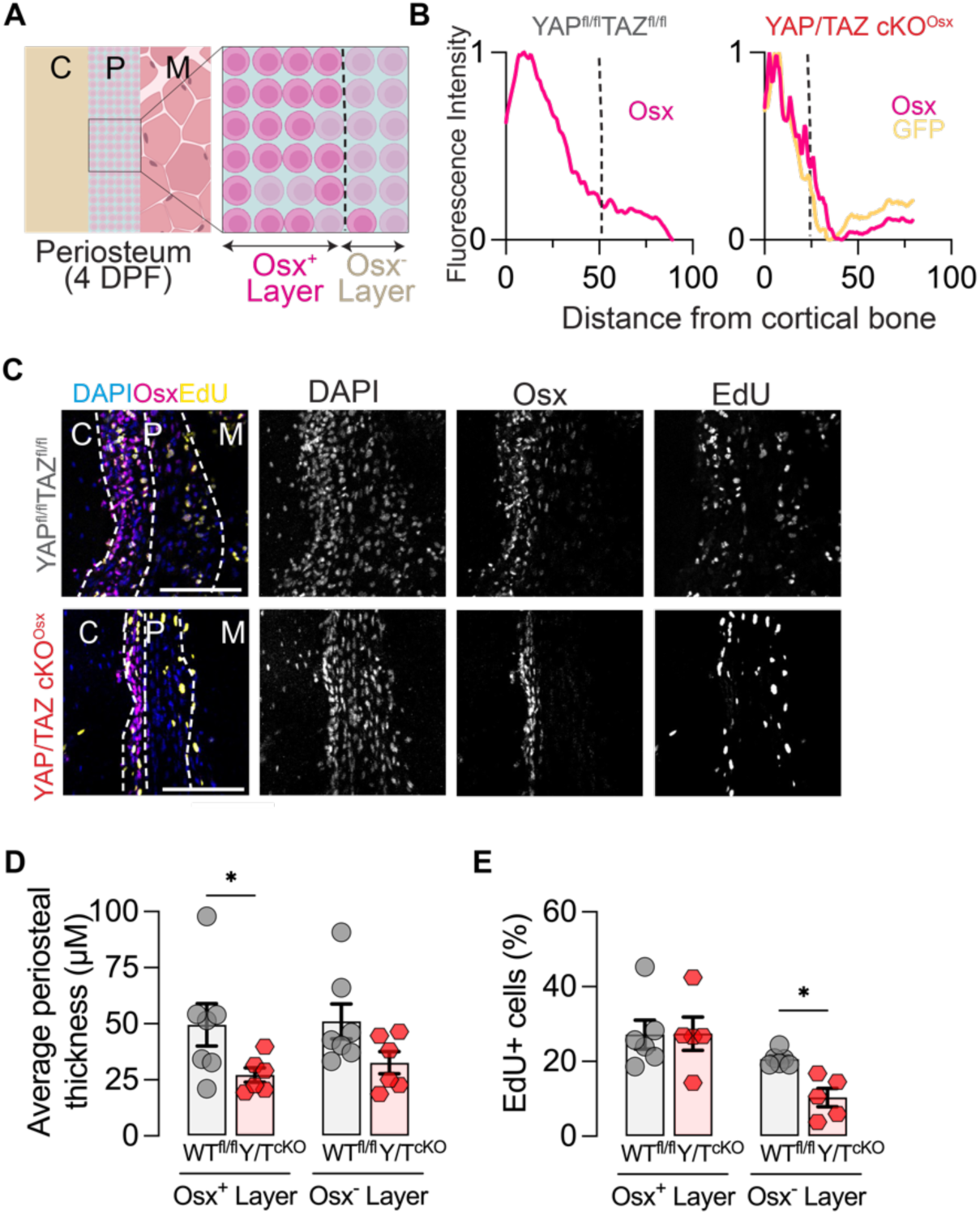
YAP/TAZ deletion from Osx-expressing cells alters both cell intrinsic and extrinsic pathways. **A.** Schematic demonstrating spatial distribution of Osx^+^ and Osx^−^ layers in the periosteum. **B.** Quantification of fluorescence intensity of Osx and Cre:GFP in WT^fl/fl^ and YAP/TAZcKO^Osx^ mice. **C.** Staining for Osx protein, EdU and DAPI. Dotted lines indicate the periosteum and the separation between the inner Osx^+^ layer and the outer Osx^−^ layer. Quantification of **D.** thickness of the Osx^+^ and Osx^−^ layers and **E.** Percentage of proliferative EdU^+^ cells. ‘*’:p<0.05.

YAP/TAZ deletion significantly decreased the average thickness of the inner Osx^+^ layer. The thickness of the outer Osx^−^ layer was also decreased, but did not reach statistical significance (**Fig. 4 D**). YAP/TAZ deletion significantly reduced cell proliferation in the outer Osx^−^ layer, while the differences in EdU labeling in the Osx^+^ layer were not significant (**Fig. 4 E**).

Next, we evaluated the number of proliferating cells. Interestingly, YAP/TAZ deletion did not reduce the number of proliferating Osx^+^ cells but significantly reduced the proliferating Osx^−^ cells (**Suppl. Fig. 1 E**). This suggests that YAP signaling in Osx^+^ cells promotes the expression of cell-extrinsic factors that may stimulate proliferation in Osx^−^ cells.

### YAP and TEAD transcribe BMP4 in periosteal cells

Since YAP-mediated transcription also depends on the DNA-binding transcription factors it associates with, we sought to identify its potential binding partners. First, we evaluated the changes in chromatin accessibility by sequencing transposase accessible chromatin sites upon YAP^CA^ expression (ATAC-Seq). We performed this analysis at the same timepoint at which we evaluated bulk mRNA-seq, to simultaneously map gene regulation and associated accessibility loci. YAP^CA^ expression up- and down-regulated approximately an equal number of genes. Based on this, we expected that YAP would equally modulate chromatin accessibility. In contrast, however, YAP^CA^ expression predominantly increased chromatin accessibility by opening 3427 chromatin loci, and only decreased accessibility at 206 chromatin loci (**Fig. 5 A**). Doxycycline treatment did not significantly alter chromatin accessibility in periosteal cells isolated from C57BL/6J mice (**Suppl. Fig. 1 D**).

**Figure 5:**
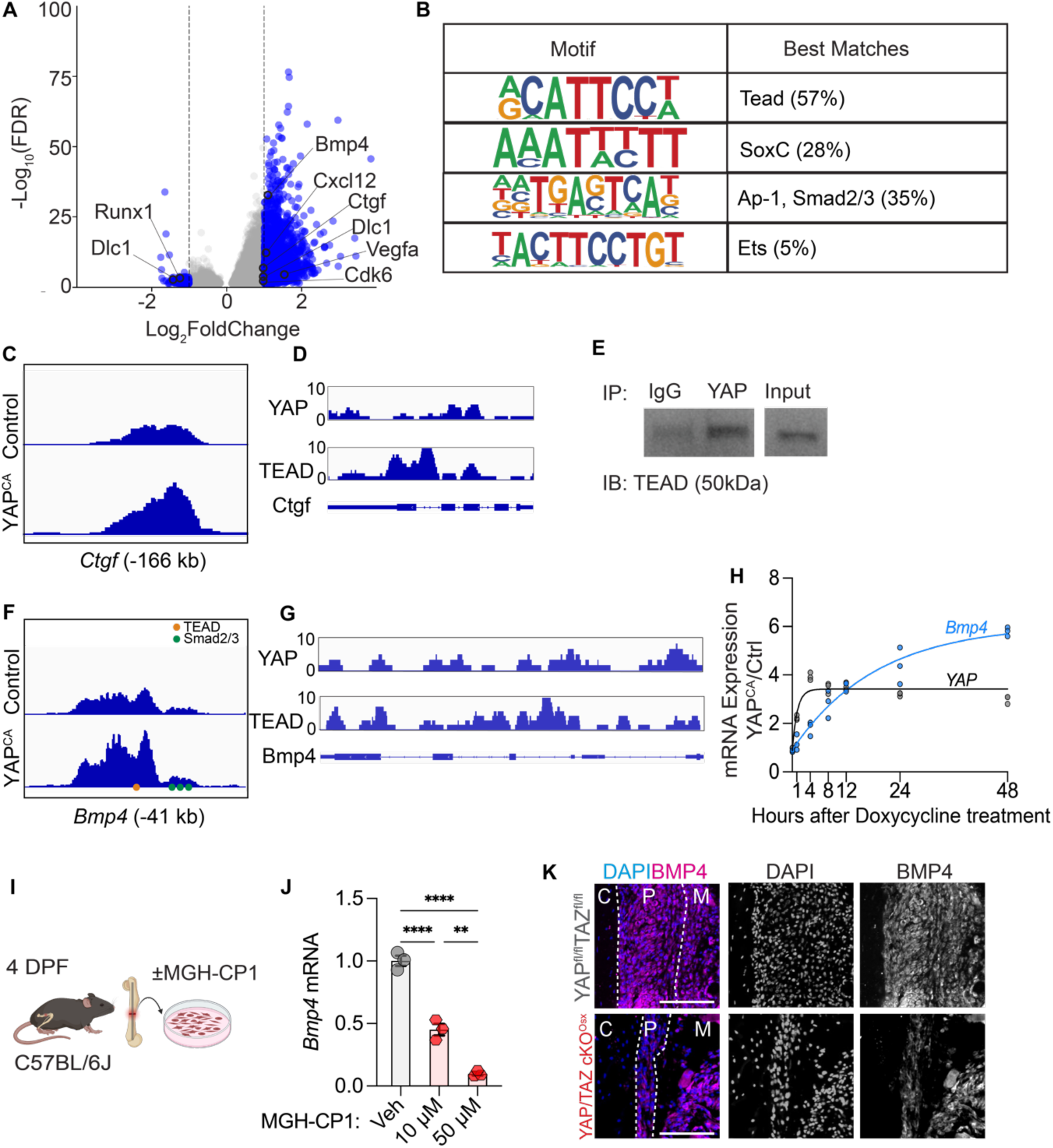
YAP transcribes BMP4 via TEAD in periosteal cells. **A.** Volcano plot of differentially accessible chromatin sites after 8 hours of YAP^CA^ expression. Log_2_FoldChange > 2 indicates more accessible chromatin after YAP^CA^. **B.** Results of motif analysis on differentially accessible chromatin loci. TEAD motifs were most frequently present, followed by those for Sox4/5/11, AP-1 cluster transcription factors and Smad2/3. **C.** Chromatin locus annotated close to *Ctgf* was more accessible after YAP^CA^ expression. **D.** YAP and TEAD pull down *CTGF* in Chromatin immunoprecipitation experiments. **E.** TEAD co-immunoprecipitates with YAP in fracture activated periosteal cells. **F.** Chromatin locus annotated close to *Bmp4* was more accessible after YAP^CA^ expression. **G.** YAP and TEAD pull down *Bmp4* in Chromatin immunoprecipitation experiments. **H.** Temporal dynamics of YAP and BMP4 expression after 1, 4, 8, 12, 24 and 48 hours of Doxycycline treatment. We observe a robust and sustained induction of Bmp4 after YAP^CA^ expression. **I.** Immunostaining for BMP4 protein in the periosteum of WT^fl/fl^ and YAP/TAZcKO^Osx^ mice. Scale bar = 100 μM. **J.** Schematic for periosteal cell isolation from C57BL6 mice followed by treatment with MGH-CP1 **K.** Treatment with MGH-CP1 results in decreased mRNA levels of *Bmp4*. ‘**’:p<0.01, ‘****’:p<0.0001.

To identify which transcription factors can bind at these chromatin loci, we performed de novo transcription factor motif enrichment analysis using Homer^15^. This unbiased analysis identified several significantly enriched sequence motifs within the differentially accessible chromatin loci. The most highly enriched motif was the consensus binding site for TEAD family transcription factors. This was followed by motifs corresponding to sox cluster genes (specifically sox4, 5 and 11), AP-1 family transcription factors and Smad2/3 (**Fig. 5 B**).

Next, we assessed chromatin accessibility near canonical YAP-TEAD target genes. YAP^CA^ expression increased chromatin accessibility at a locus proximate to *Ctgf*^5^ (**Fig 5. C**). Analysis of publicly available YAP and TEAD ChIP-seq data^16^ (<u>GSE163459</u>) indicate significant immunoprecipitation of the *Ctgf* gene locus with either YAP or TEAD pull-down (**Fig. 5 D**). To verify whether YAP interacts with TEAD in periosteal cells, we performed co-immunoprecipitation of YAP protein and probed for TEAD using a pan-TEAD antibody. TEAD family proteins were detected in the YAP pulldown, but not in the IgG control, confirming physical YAP-TEAD interaction in periosteal cells (**Fig. 5 E**).

The most differentially upregulated gene by YAP^CA^ activation was *Bmp4*. *Bmp4* mRNA is expressed in the muscle and periosteum surrounding the fracture callus^17,18^. However, it’s role in fracture repair is not well understood. Therefore, we analyzed whether YAP^CA^ increased chromatin accessibility near *Bmp4.* And found increased accessibility at a locus upstream of the *Bmp4* transcription start site (−41kb) (**Fig. 5 F**). Further, ChIP-Seq data from human endothelial cells indicates immunoprecipitation of the *BMP4* gene with both YAP and TEAD ^19^ (**Fig. 5 G**). YAP^CA^ activation dynamically induced Bmp4 mRNA expression within the first 8 hours of YAP activation (**Fig. 5 H**), which was sustained over 48 hours. Inhibition of TEAD interaction with YAP using MGH-CP1^20^, suppressed *Bmp4* expression *in vitro* (**Fig. 5 I, J**). BMP4 protein was abundant throughout the fracture-expanded periosteum and was lowered in YAP/TAZcKO^Osx^ mice (**Fig. 5 K**). Together, these data identify *Bmp4* as a YAP-TEAD target gene that may be involved in the periosteal response to fracture.

### BMP4 delivery improves periosteal expansion in the outer layer of the periosteum

Therefore, we tested whether exogenous BMP4 treatment could improve periosteal expansion in YAP/TAZ cKO^Osx^ mice. To this end, we injected recombinant mouse BMP4 protein in the fracture gap of YAP/TAZ cKO^Osx^ mice daily until 4 DPF (**Fig. 6 A**). BMP4 treatment increased periosteal thickness (**Fig. 6 C**) to levels comparable to what we observe in WT^fl/fl^ mice post fracture (shaded area). However, BMP4 treatment did not affect overall cell proliferation (**Fig. 6 D**), nor did it alter the total numbers of Osx^+^ or Osx^−^ cells (**Suppl. Fig. 2 A, B**). Next, we looked at the differences between the inner Osx^+^ and outer Osx^−^ layers. Interestingly, the increase in periosteal thickness after BMP4 treatment was driven by the expansion of the outer Osx^−^ layer. (**Fig. 6 E**). Additionally, cell proliferation also increased in the Osx^−^ layer after BMP4 treatment (**Fig. 6 F**).

**Figure 6:**
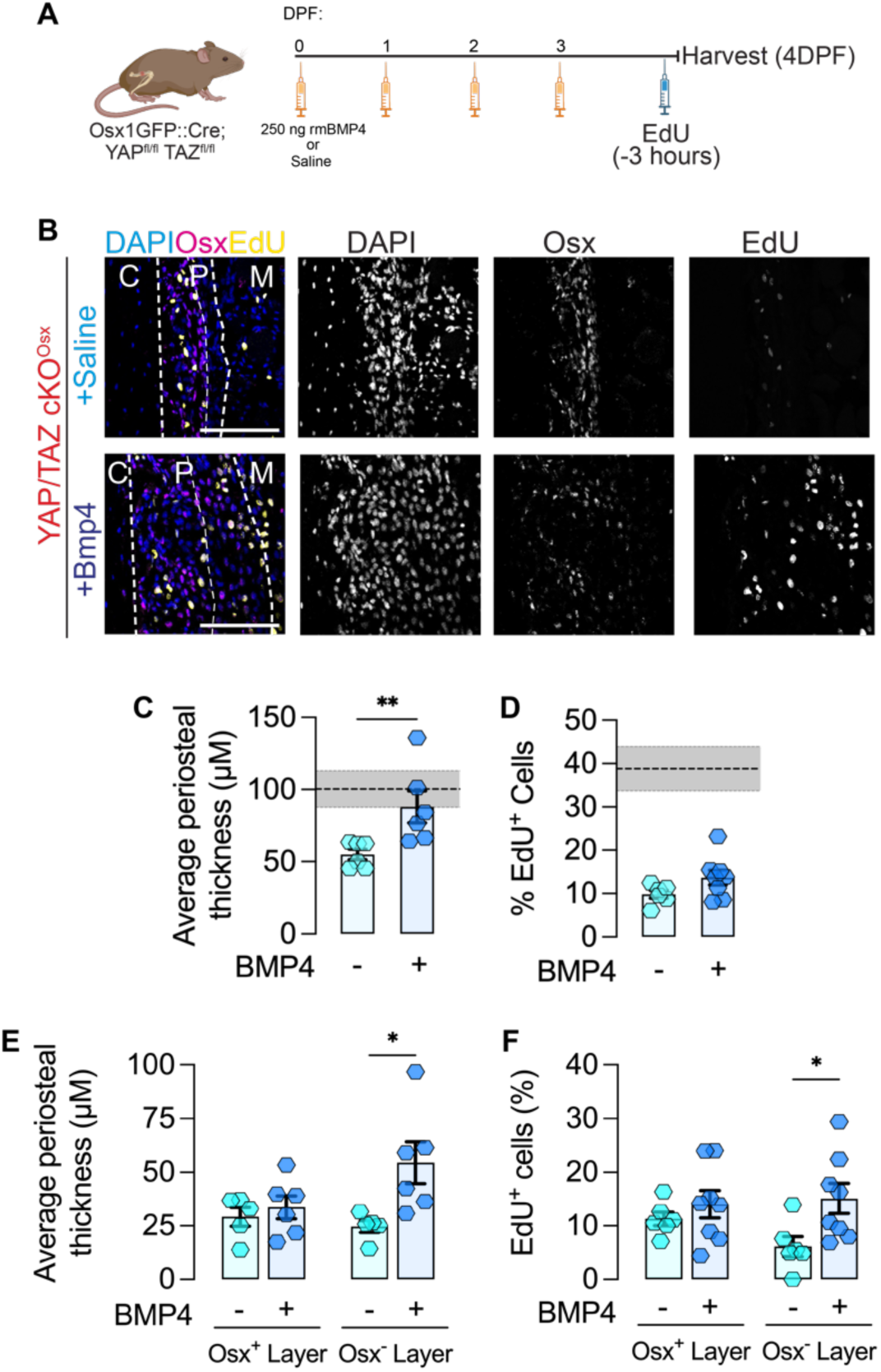
*In vivo* delivery of Bmp4 during early stages of fracture repair rescues periosteal expansion in the outer Osx- layer. **A.** Timeline of *in vivo* rmBMP4 delivery for 4 days after femoral osteotomy. **B.** Immunofluorescence staining of Osx with DAPI (nuclei) and EdU (proliferating cells) in the inner and outer layers of the periosteum in YAP/TAZcKO^Osx^ mice injected with saline (top panel) or BMP4 (bottom panel). **C.** BMP4 treatment increases average periosteal thickness in YAP/TAZcKO^Osx^ mice. **D.** Percentage of EdU^+^ cells remain unchanged. Gray bar indicates the range of values in injured 4DPF WT^fl/fl^ mice. Quantification of **E.** thickness of the Osx^+^ and Osx^−^ layers and **F.** percentage of proliferative EdU^+^ cells in each layer. Scale bar = 100 μM. ‘*’:p<0.05.

To further explore the mechanism behind BMP4-mediated periosteal expansion, we examined chondrogenic and osteogenic activity in the periosteum. First, we looked for Sox9^+^ chondroprogenitors in the periosteum. A majority of the Sox9^+^ cells were also positive for Osx, in both WT^fl/fl^ and YAP/TAZcKO^Osx^ mice, irrespective of BMP4 treatment (**Suppl. Fig. 3 A-F**). Based on our data and previous findings by other groups^21^, these cells are likely osteo-chondroprogenitors that have not yet committed to an osteoblast or chondrocyte lineage at 4 DPF. YAP/TAZ deletion or BMP4 treatment in knockout mice did not affect chondrogenesis (as indicated by Sox9 staining and GAG staining (**Suppl. Fig 3 A, D**) or ALP activity (**Suppl. Fig. 4 A-F**).

Since YAP/TAZ signaling and BMP4 both play a role in angiogenesis^22,23^, we examined blood vessels in the periosteum by staining for Endomucin (Emcn) (**Suppl. Fig. 5 A, E**). YAP/TAZ deletion and BMP4 treatment did not change Emcn^+^ area (**Suppl. Fig. 5 B, F**), vessel area fraction (**Suppl. Fig. 5 C, G**) or the size of blood vessel loops (**Suppl. Fig. 5 D, H**) in the periosteum.

### BMP4 delivery promotes periosteal expansion by collagenous matrix production

Given that YAP activation altered expression of genes and gene sets associated with matrix organization, we asked whether the exogenous BMP4 administration affected the ECM in the periosteum. Treatment of fracture-activated periosteal cells *in vitro* with recombinant mouse BMP4 upregulated BMP4 target genes *Id1* and *Serpine1* within 24 hours (**Fig. 7 A, B, C**). By day 4, *Col1a1* mRNA levels were elevated (**Fig. 7 D**). Finally, we used second harmonic generated (SHG) microscopy to assess the collagen matrix in the expanding periosteum (**Fig. 7 E, F**, **Suppl. Fig. 6 A, B**). SHG signal was significantly higher in the periosteum after BMP4 treatment in YAP/TAZ cKO^Osx^ mice. Together, these data show that YAP transcribes BMP4 via TEAD, and BMP4 contributes to periosteal expansion by increasing matrix deposition.

**Figure 7:**
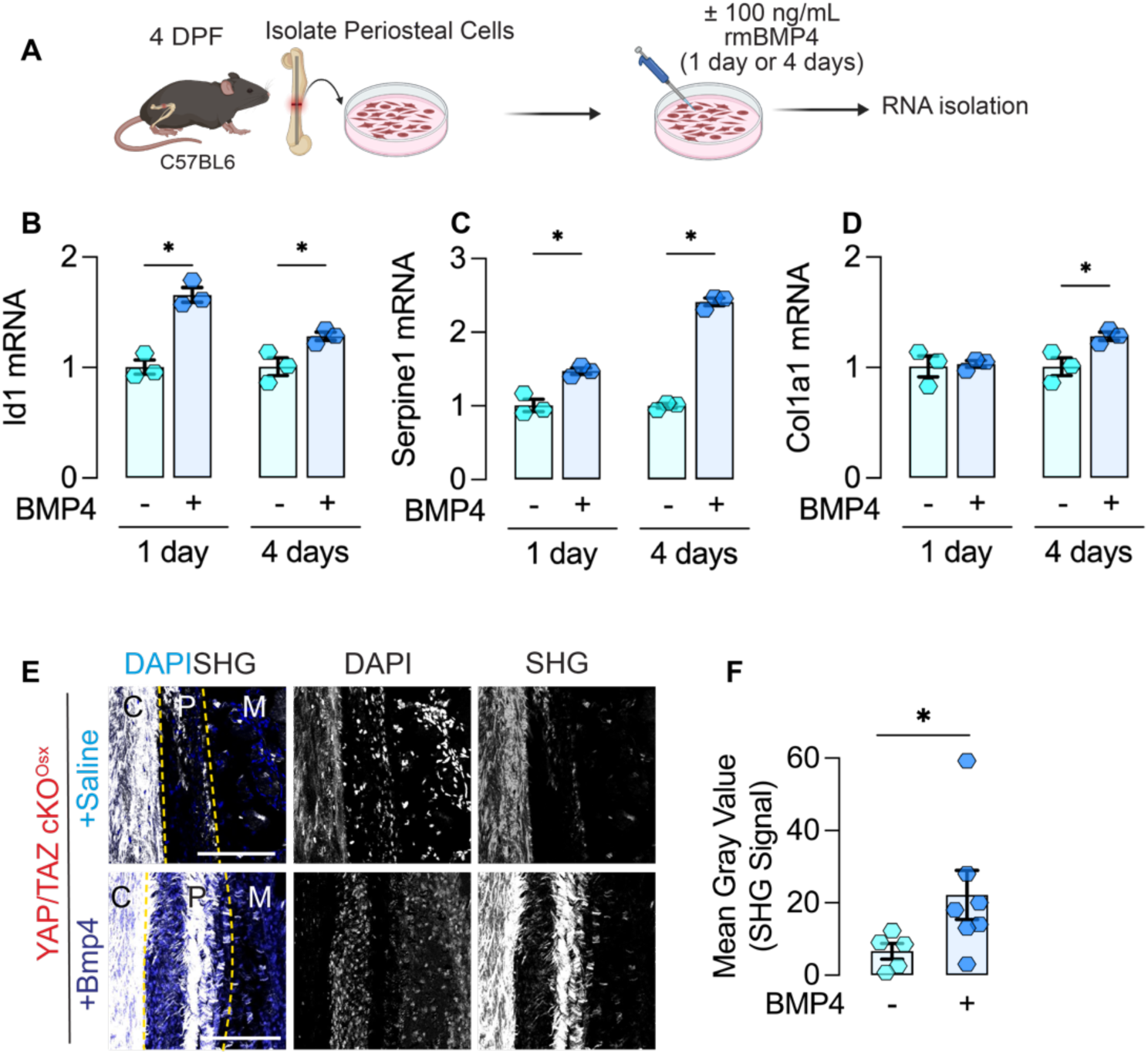
BMP4 promotes collagen matrix production in the periosteum. **A.** Schematic for periosteal cell isolation from C57BL6 mice at 4DPF, followed by rmBMP4 treatment *in vitro* for 1 or 4 days. BMP4 treatment results in a robust induction of mRNA levels of **B.** *Id1* **C.** and *Serpine1* both 1- and 4-days post treatment. **D.** *Col1a1* mRNA transcripts are significantly higher after 4 days of BMP4 treatment compared to saline treated controls. **E.** SHG imaging of periostea in YAP/TAZcKO^Osx^ mice after BMP4 treatment shows increased collagen matrix deposition. **F.** Quantification of SHG intensity in YAP/TAZcKO^Osx^ mice after BMP4 treatment. Scale bar = 100 μM. ‘*’:p<0.05.

## Discussion

Here we demonstrate that YAP is activated in periosteal cells after fracture and regulates transcriptional programs that promote periosteal expansion via both cell-intrinsic and cell-extrinsic factors. Conditional deletion of YAP and TAZ from Osterix-expressing cells, which reside in the inner layer of the periosteum, impaired expansion of both the Osx^+^ inner layer and the Osx^−^ outer layer. We identified BMP4 as a YAP-TEAD target gene induced by YAP activation. Exogenous BMP4 delivery improved periosteal expansion in YAP/TAZ cKO mice and enhanced periosteal expansion through cell proliferation and matrix deposition. Together, these data uncover new insights into the transcriptional mechanisms by which YAP regulates the early stages of fracture repair.

YAP regulates transcriptional programs that promote periosteal expansion via both cell-intrinsic and cell-extrinsic mechanisms. We refer to cell-intrinsic factors as genes that can regulate the behavior of the same cell, and cell-extrinsic factors as genes that can regulate the behavior of neighboring cells. We designed this study to capture the earliest YAP target genes to identify direct YAP targets via RNA- and ATAC-seq. We found that even at the early time point, YAP induces changes to both cell-intrinsic and cell-extrinsic programs as indicated by Gene Set Variation Analysis. Direct YAP target genes reflect these intrinsic and extrinsic functions. For example, YAP regulates the cell-intrinsic factor, *Cdk6*. *Cdk6* encodes a Cyclin Dependent Kinase that regulates cell proliferation^24^. Similarly, YAP regulates the cell-extrinsic factor *Il6,* which encodes an interleukin that regulates immune cell recruitment after fracture^25^. While we observed similar numbers of up- and down-regulated genes by RNA-Seq, YAP activation mostly increased chromatin accessibility as assessed by ATAC-Seq. This indicates that while YAP enhances chromatin accessibility, it can act as both a transcriptional activator and repressor in these cells.

*In vivo,* YAP deletion from Osx^+^ cells influenced Osx^+^ and Osx^−^ cells. This is consistent with the idea that YAP regulates both cell-intrinsic and cell-extrinsic signaling. We also observed changes in the Osx^+^ and Osx^−^ layers of the periosteum, which suggests a YAP-mediated regulation of cell-cell communication between the two layers. Notably, YAP activation induced a robust increase in transcript levels of *Bmp4*, which is an established cell-extrinsic growth factor – and could be a mediator of cell-extrinsic signaling in the early stages of fracture repair.

In this study, we demonstrated that YAP-TEAD can drive *Bmp4* transcription in periosteal cells. Previous studies from other groups have shown that BMP4 mRNA is present in the proliferating periosteum around the fracture gap^18^, supporting the idea that periosteal cells are a source of BMP4. Together, these data suggest that periosteal cells secrete BMP4. While BMPs play a key role in bone development and repair^26^, the specific function of BMP4 in bone regeneration remains unclear^18,27^. At the transcriptional level, we confirmed that YAP and TEAD together transcribe *Bmp4.* The preferentially open BMP4 locus also contains motifs for Smad2/3, and BMP4 is known to regulate its own expression through feedback and feedforward loops involving SMAD transcription factors^28^. Further, Smad2/3 can interact with co-factors like YAP and TAZ to regulate gene expression in other contexts of tissue repair and development^29^.

Exogenous BMP4 treatment improves periosteal expansion in mice lacking YAP/TAZ in Osx^+^ cells. This effect was driven by the expansion of the outer layer of the periosteum, where we observed increased matrix deposition and cell proliferation. Increase in cell proliferation was restricted to the outer layer of the periosteum, whereas matrix deposition was elevated across the periosteum. This distinction may be due to the delivery method we used, or because of cell-specific responsiveness to BMP4 in the outer layer. Previous studies with other BMPs suggest that the delivery method can significantly impact outcomes^30,31^. We did not observe altered osteogenesis, angiogenesis or chondrogenesis after BMP4 treatment. However, using a sustained delivery method – such as slow-release hydrogels or collagen sponges – could potentially yield different results. Further research is needed to elucidate the role of BMP4 and its downstream pathways in periosteal expansion and fracture repair.

Together, these data demonstrate that YAP regulates periosteal expansion during early fracture repair by driving cell-intrinsic and extrinsic transcriptional programs, including the TEAD-mediated transcription of BMP4, which promotes proliferation and matrix production in the outer periosteal layer.

### Limitations

This study has several limitations. First, while YAP and TAZ combinatorially regulate bone development and regeneration^4,32^, our gain-of-function approach targeted only YAP and did not address the co-transcriptional roles for TAZ in periosteal cells. While our loss of function and rescue experiments featured both YAP and TAZ deletion, further research will be required to identify the specific roles of TAZ in periosteal gene regulation. Second, our study focused on BMP4 as a YAP target gene, but YAP regulates a diversity of gene targets in fracture-activated periosteal cells, which we have not explored. Although exogenous BMP4 improves periosteal expansion, it is certainly not the only contributing factor. Third, while this study identifies factors (including BMP4) that could be introduced exogenously, this study was not designed to evaluate their therapeutic potential. Additional research, including long-term assessment of fracture repair efficiency following exogenous treatment, is needed to determine whether these factors could be effectively used as therapeutics. Lastly, while YAP and TAZ deletion in Osx^+^ cells impacted both Osx^+^ and Osx^−^ populations, it remains unclear whether Osx^−^ cells originate from the Osx^+^ population and, if so, whether YAP/TAZ were also deleted in the Osx^−^ cells. Although we did not perform lineage tracing, prior studies indicate that cells from the Osx^−^ and Osx^+^ layers maintain distinct identity throughout fracture repair^14^.

## Materials and Methods

### Animals

#### Osterix conditional YAP/TAZ knockout mice

Osx-Cre::GFP;YAP^fl/fl^TAZ^fl/fl^ Mice harboring loxP-flanked exon 3 alleles in both YAP and TAZ on a mixed C57BL/6J genetic background were provided by Dr. Eric Olson (University of Texas Southwestern Medical Center). Tetracycline-responsive B6.Cg-Tg(Sp/7-tTA,tetO-EGFP/Cre)1AMc/J (Osx-Cre^tetOff^) mice from The Jackson Laboratory (Bar Harbor, ME, USA) were used to generate a mouse model in which we conditionally deleted YAP and TAZ from Osx expressing cells. In this mouse model, tetracycline (or its more stable derivative, doxycycline) administration prevents tetracycline-controlled transactivator protein (tTA) binding to the tetracycline-responsive promoter element (TRE) in the promoter of the Cre transgene, allowing Cre expression only in the absence of doxycycline for temporal control of Osx-Cre mediated gene deletion.

We allowed mice to develop to skeletal maturity and induced homozygous YAP/TAZ deletion 2 weeks before fracture between 14-16 weeks of age. All mice were bred and raised until skeletal maturity with doxycycline in their drinking water to prevent Cre-mediated gene recombination. For all *in vivo* fracture healing assessments, doxycycline was withdrawn from drinking water 2 weeks before fracture surgery and normal drinking water was provided for the remainder of the study. We used YAP^fl/fl^TAZ^fl/fl^ mice (WT^fl/fl^), without the Osx-Cre:GFP as controls. We have previously shown that these are comparable with Osx-Cre:GFP mice in bone development and fracture repair^4,23^.

#### YAP^S127A^ mice for gain-of-function experiments

We received mice harboring tet inducible YAP^S127A^ allele tetO-YAP^S127A^;Gt(ROSA)26^Sortm1(rtTA,EGFP)Nagy/J^ from Dr Jenna Galloway (Massachusetts General Hospital/Harvard Medical School). These were bred with mice harboring a CMV-Cre allele (CMV-Cre B6.C-tg(CMV-Cre)1Cgn/J, Jackson Laboratory, Bar Harbor, ME, USA). Femoral bone fractures were induced surgically, and cells were isolated from these mice at 4 DPF. Cells were treated with doxycycline to induce YAP^S127A^ or YAP^CA^ expression. Untreated cells were used as controls.

All mice were ear clipped after weaning and genotyped through an external service (Transnetyx Inc., Cordova, TN, USA). All mice were fed regular chow (PicoLab Rodent Diet, Cat. 0007688, Purina LabDiet, St. Louis, MO, USA) *ad libitum* and housed in cages containing 2-4 animals each. Mice were maintained at constant 25°C on a 12-hour light/dark cycle. Both male and female mice were evaluated with the same fracture healing procedure. All surgeries were performed on mice between 14-16 weeks of age. All protocols were approved by the Institutional Animal Care and Use Committees at the University of Pennsylvania (Protocol no: 806482). All animal procedures were performed in adherence to federal guidelines of animal care and following the 3R principles in animal research^33^.

### Femoral fracture surgeries and time points

An open, unilateral, intramedullary pin-stabilized femoral fracture model was used to study bone repair. Animals were anesthetized using isoflurane (1-3%). All hair was removed from the surgical site using a hair trimmer, and the area was cleaned with betadine and 70% Ethanol. Buprenorphine-SR (Covetrus, Portland, ME, USA) was applied subcutaneously at 1mg/kg before transferring mice to the sterile surgical field. Femora were accessed through a lateral incision made on the right hindlimb followed by blunt dissection of muscles. Femoral condyles were visualized, and a 25-gauge needle was inserted from between the condyles to stabilize the bone. Muscle surrounding the femur was blunt dissected to expose the femoral midshaft. Osteotomy was performed using a 0.45mm Gigli wire saw (RISystem, 590.110, Landquart, Switzerland). The incision site was closed using nylon sutures. Mice were allowed to recover under a heating lamp and transferred to regular cages after complete recovery. To ensure food and water intake after surgery DietGel (ClearH_2_O, Westbrook, ME, USA) was provided on the cage floor. Humane endpoints were predefined, including criteria such as wound dehiscence, poor grooming, sunken eyes, hunched posture, severe axial deviation, lack of food and water intake, significant weight loss, bloody feces, severe respiratory issues, debilitating diarrhea, seizures, paresis, and abscesses. No humane endpoints were reached during the study. Mice were monitored closely until experimental endpoints (4DPF). 3 hours prior to euthanasia, mice were injected intraperitoneally with 4-ethynyl-2’-deoxyuridine (EdU; E10187; Invitrogen, Waltham, MA, USA) at 10mg/kg to assay cellular proliferation. Animals were euthanized by CO_2_ inhalation followed by cervical dislocation.

### Cryohistology and Immunofluorescence staining

Bones harvested at 4DPF were fixed in 4% Paraformaldehyde (PFA) at room temperature for 4 hours, followed by decalcification in 20% ethylenediaminetetraacetic acid (EDTA) for 72 hours. Decalcified bones were transferred to 30% Surcose in 1X Dulbecco’s phosphate buffered saline (DPBS) for 48 hours and embedded in optimal cutting temperature (OCT) Compound (Tissue-Tek, Torrance, CA, USA). 10 μM thick tissue sections were obtained on a cryotape (Section Lab Co, Hiroshima, Japan) as described in a previously published protocol^34^. Sections were stored at −20°C until they were ready for staining. Tape sections were glued to microscope slides with Norland Optical Adhesive 81 (Norland Products Inc, Jamesburg, NJ, USA), blocked with 5% goat serum in 0.3% Triton-X-100. Wherever required, EdU was detected using the Click-iT EdU detection kit (C10337 or C10339, Invitrogen, Carlsbad, CA, USA). This was followed by primary and secondary antibody staining. A list of antibodies used in this study is provided in the supplementary material. Samples were imaged using a Zeiss AxioScan or a Zeiss LSM 710 Confocal microscope (Carl Zeiss Microscopy, Thornwood, NY).

### Periosteal cell isolation and *in vitro* cell culture

Mouse periosteal cells were isolated from either C57BL/6J or YAP^CA^ mice from 4-day old fracture calluses. Briefly, mice were euthanized by CO_2_ inhalation followed by cervical dislocation. All non-osseus tissue was dissected off the fractured femora. The epiphyses were removed, and marrow cavities were flushed. The periosteum was scraped and enzymatically digested for 1 hour at 37°C on an orbital shaker (0.5 mg/mL collagenase P, 2 mg/mL hyaluronidase in PBS). Digestion was stopped by adding complete growth medium (**α**-MEM, 15% FBS, 1% Penicillin-Streptomycin, 1% Gluta-MAX I, 0.1% **β**-Mercaptoethanol). Digested tissue and cells were passed through a 70 μM cell strainer to remove debris. The resulting cell suspension was centrifuged at 300 x g for 5 minutes, re-suspended in complete growth medium and plated on cell culture flasks (1 limb per 25cm^2^ flask) at 5% O_2_. After 3 days, flasks were transferred to 20% O_2_ for subsequent processing. All *in vitro* analyses were performed on cells passaged 1-5 times. For all experiments, cells were passaged at 5000 cells/cm^2^ on tissue culture plates. Cells were switched to 10% serum during drug or growth factor treatments.

### mRNA isolation for qPCR and bulk mRNA-sequencing

RNA was isolated from cultured cells using the RNeasy Mini Kit (74106, Qiagen, Hilden, Germany), and quantified using a NanoDrop 2000 (ThermoFisher Scientific, Waltham, MA, USA). 0.5μg of RNA from each sample was used to make complementary DNA using a High-Capacity cDNA Reverse Transcription Kit (4368814, Thermo-Fisher Scientific). Quantitative polymerase chain reaction (qPCR) was performed using QuantStudio Pro 6 Real-Time PCR System (Thermo-Fisher Scientific, Waltham, MA). Relative quantification of target genes was calculated by the ΔΔCt method with *Hprt* and/or *18S rRNA* as internal controls. For all *in vitro* studies, each data point for qPCR represents a separate well on a cell culture plate.

### Bulk mRNA-sequencing and analysis

RNA isolation was performed as described in the section above. Paired end RNA sequencing data was aligned to the mm10 genome assembly using hisat2 and uniquely mapped reads were counted using featureCounts^35^ with mm10 GTF file from NCBI Ref Seq. We used DESeq2^36^ v1.32.0 in RStudio (2022) using a fold change threshold of 0.7<FC<1.3 and a p value of 0.05. Fold change thresholds were decided based on the level of upregulation of canonical YAP target genes *Ctgf* and *Cyr61*. Data visualization was performed using packages included in DESeq2 and EnhancedVolcano (Blighe, Rana and Lewis 2023, Bioconductor).

### ATAC-sequencing and analysis

50,000 cells per technical replicate were subjected to Assay for Transposase Accessible Chromatin Sequencing (ATACSeq) following a published protocol^37^. Paired-end reads were mapped to the mm10 genome using bowtie2^38^. Picard (https://broadinstitute.github.io/picard/) SortSam was used to convert SAM files to BAM files. PCR duplicates were removed using Picard MarkDuplicates. Mitochondrial reads were removed after indexing BAM files with SAMtools^39^. Processed BAM files were used to call peaks using MACS2^40^. Peaks in the blacklisted regions of the genome were discarded from subsequent analysis using SAMtools as described previously^41,42^. Diffbind (Stark and Brown, 2011, Bioconductor) was used as implemented in the Galaxy platform to identify differentially accessible chromatin loci between groups. Peaks were visualized using Integrative Genomics Viewer (IGV, Broad Institute, MA, USA). Bedtools was used to generate bigwig files suitable for IGV. Motif analysis was completed using the findMootifsGenome.pl command in Homer.

### Gene Set Variation Analysis

To uncover the functional changes demonstrated by transcriptomic differences, gene set variation analysis (GSVA) was performed using the transcript counts. This analysis was conducted with the R GSVA package (v1.52.1) downloaded from Bioconductor. The gene sets used were the human “C2: curated gene sets” from MSigDB. The minimum size for gene sets was set at 15 transcripts and all others were removed. To establish differential enrichment, permutation testing was utilized, as previously described. To perform this testing, the true mean enrichment difference between groups was determined using the GSVA function. The gene names in the count matrices were then randomly shuffled, and the mean enrichment difference was recalculated. This random shuffling was repeated for a total of 1000 times. The p-value for this analysis was determined as the proportion of times that the random shuffling produced a mean difference that was greater in magnitude than the true difference. Benjamini and Hochberg false discovery rate adjustment was used for adjustment. An FDR adjusted p-value less than 0.05 was considered significant.

### Protein Lysis and Western Blotting

*In vitro* cells were lysed in RIPA Buffer with protease and phosphatase inhibitors. Protein quantification was performed using a Pierce BCA Kit (Cat. 23227, Thermofisher Scientific, Waltham, MA, USA). Samples were normalized for protein and run on a Bio-rad MINI Protean gel in 1X Tris/Glycine/SDS running buffer. Blot was transferred onto a PVDF membrane using a BioRad Turbo Transfer device. Proteins were detected following manufacturer described protocols for specific antibodies. Each experiment was repeated 2 times with fresh lysates.

### Coimmunoprecipitation

Proteins were lysed using a Cell Lysis buffer (Cat. 9803, Cell Signaling Technology, Danver, MA, USA) with PMSF (protease inhibitor) and a phosphatase inhibitor (PhosSTOP tablets). Protein quantification was performed using a BCA kit. All lysates were at a concentration of 1mg/mL. Primary antibodies (either IgG or YAP) were concentration matched and added to protein lysates and incubated overnight at 4°C. Antibody and protein complex was pulled down using Protein A and G Magnetic beads and run on a BioRAD mini protean gel as described above. Blots were transferred to PVDF membranes and probed for target proteins with antibodies. As the proteins we were probing for were all in the range of 50-60kDa, we used a light chain specific, HRP conjugated secondary antibody for detection (Cat. 45262, Cell Signaling Technology).

### Analysis of published ChIP-Seq data

Publicly available fastq files were obtained from Gene Expression Omnibus (GSE163458)^19^. Fastq files were aligned to the GRCh38 build using bowtie2 with default settings^38^. Resulting aligned SAM files were converted to BAM files and sorted by chromosomal coordinates using Picard SortSam. Mitochondrial DNA and duplicates were removed with Picard. MACS2 was used to call peaks using the Input as the background with an FDR q-value cutoff of 0.01^40^. Input-subtracted bigWig files were generated with bedtools, genomeCoverageBed, and bedGraphtoBigWig tools^43,44^. Resulting bigWig tracks were visualized using Integrative Genome Viewer on the hg38 genome track.

### Statistical analysis and figure preparation

All graphs report Mean and SEM. *In vivo* studies: Sample sizes were selected based on power analyses previously performed for our mouse models^4^. Comparisons between two groups were made using unpaired t-tests. When parametric test assumptions were not met, a Mann-Whitney test was used. A p-value<0.05 was considered significant. For *in vitro* sequencing experiments, periosteal cells were pooled from 3 mice of the same genotype and plated in 3 different wells as technical replicates. Statistical analyses for sequencing experiments are included in their respective methods sections. All analyses were performed blind to the groups (genotype, treatment), and de-blinded only after completion to avoid bias.

### Antibodies

| Protein detected | Manufacturer | Catalog Number | Concentration | Application |
| --- | --- | --- | --- | --- |
| YAP | Cell Signaling Technology | 14074 | 1:100 | In vivo IF |
| Human YAP | Santa Cruz Biotechnology | sc-376830 | 1:100 | In vitro IF |
| Osterix/Sp7 | Abcam | ab209484 | 1:100 | In vivo IF |
| Sox9 | Abcam | ab196184 | 1:100 | In vivo IF |
| Endomucin | Santa Cruz Biotechnology | sc-65495 | 1:100 | In vivo IF |
| BMP4 | Abcam | ab39973 | 1:100 | In vivo IF |
| YAP/TAZ | Cell Signaling Technology | 8418 | 1:1000 | Western Blot |
| β-Actin | Cell Signaling Technology | 4967 | 1:1000 | Western Blot |

### Primers

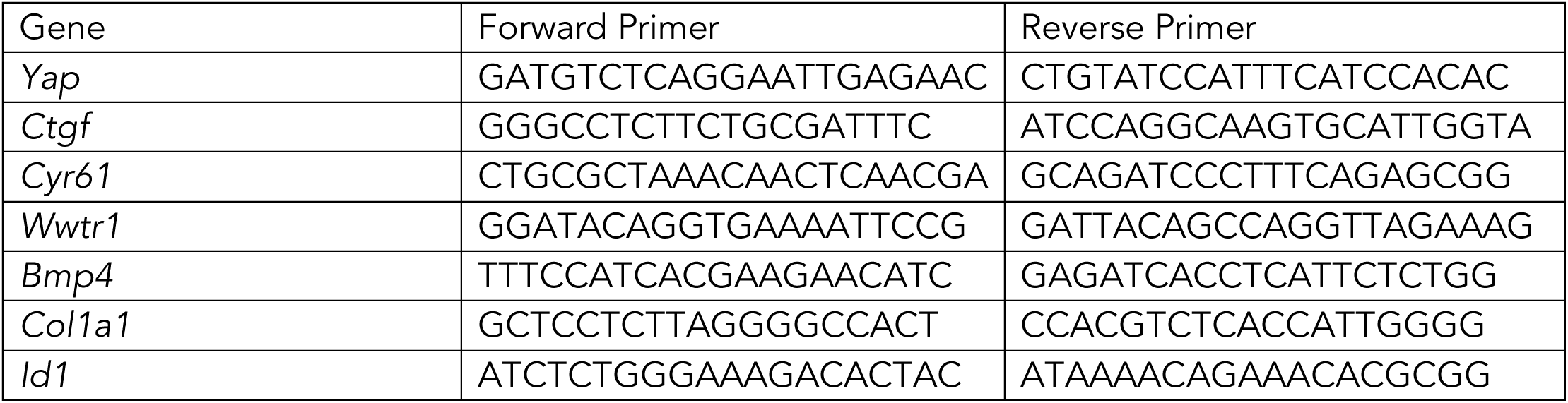
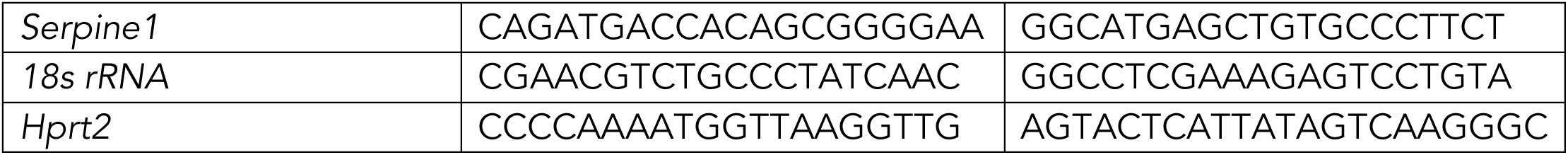

## Acknowledgements

The authors would like to thank all members of McKay Orthopedics Research Lab for constructive inputs and discussion.

## Funding Sources

This study was funded by NIH/NIAMS R01-AR074948, R01-AR073809, P30-AR050950, NSF/CMMI:15-48571 (to J.D.B), NIH/NIGMS K12GM081259 (to C.J.P.), NIH/NRSA T32AR007132 (to E.S.). A.L. received a Research Fellowship funded by the Deutsche Forschungsgemeinschaft (DFG; project no.: 440525257; reference: LA 4007/2-1).

## Author Contributions

M.P.N. and J.D.B conceived and supervised the research. M.P.N., A.L., M.B., G.I.T., C.J.P., E.S. and Y.M. performed experiments. M.P.N., B.T., D.L.J., G.H., L.W., analyzed data. G.L.S., and N.A.D., supported with methods and technical advice. M.P.N. and J.D.B. wrote the paper. All authors discussed and revised the manuscript.

## Competing interests

G.L.S. is an employee of, and holds equity in, Pfizer. The authors declare that they have no competing interests.

## Data Availability

All data are available in the main text or supplementary materials. The data discussed in this publication have been deposited in NCBI’s Gene Expression Omnibus and are accessible through GEO Series accession number GSE280239.

---

**Supplementary Figure 1:**
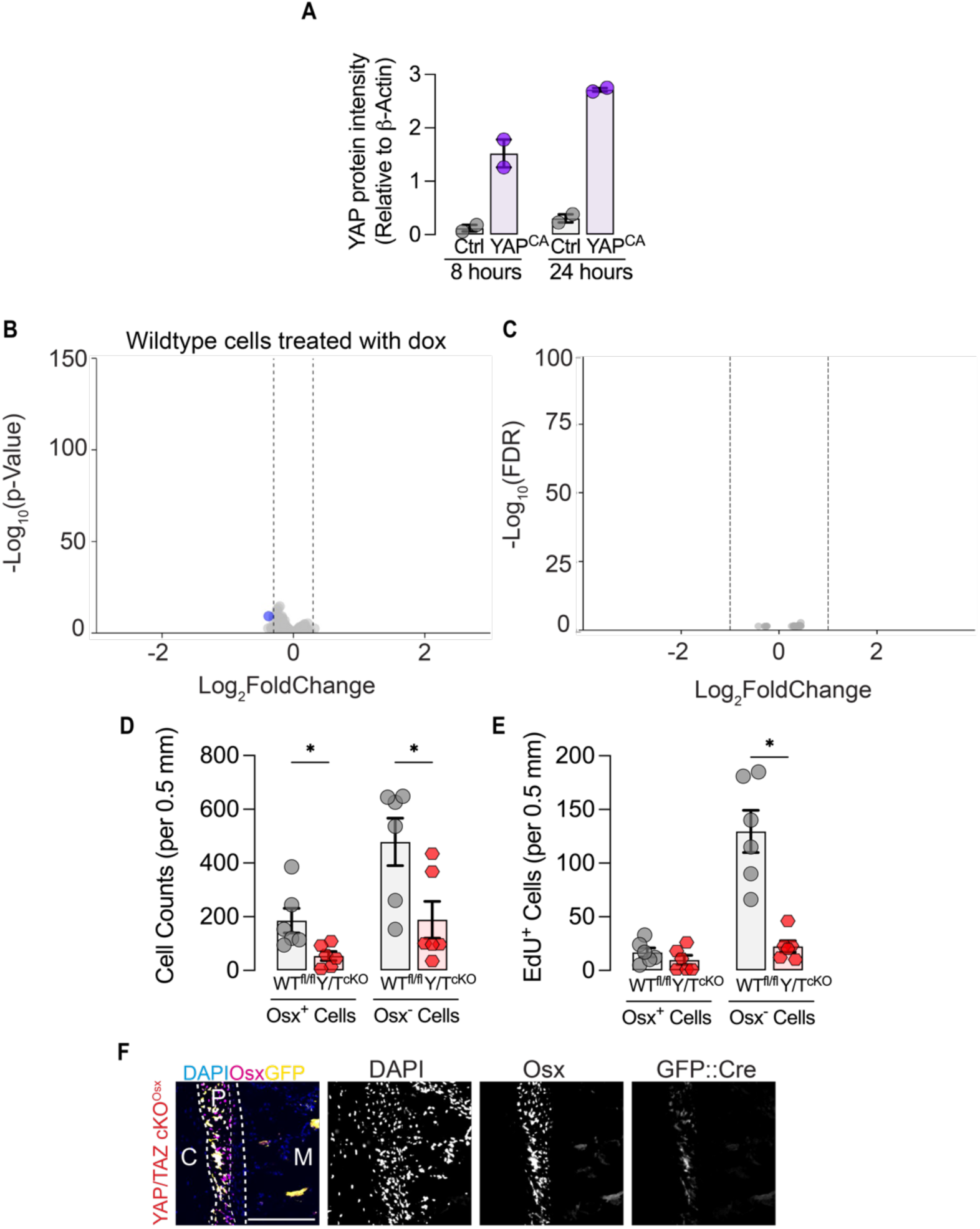
**A.** Quantification of protein intensity of YAP (relative to β-actin) after 8 and 24 hours of doxycycline treatment. **B.** 8 hours of doxycycline treatment does not affect gene expression in periosteal cells derived from C57BL6 mice. **C.** 8 hours of doxycycline treatment does not affect chromatin accessibility in periosteal cells derived from C57BL6 mice as assessed by ATAC-Seq. **D.** Total counts of Osx^+^ and Osx^−^ cells in the periosteum in WT^fl/fl^ and YAP/TAZcKO^Osx^ mice. **E.** Total counts of proliferating Osx^+^ and Osx^−^ cells in the periosteum in WT^fl/fl^ and YAP/TAZcKO^Osx^ mice. **F.** Immunostaining for Osx and GFP in the periostea of YAP/TAZcKO^Osx^ mice. Scale bar = 100 μM.

**Supplementary Figure 2:**
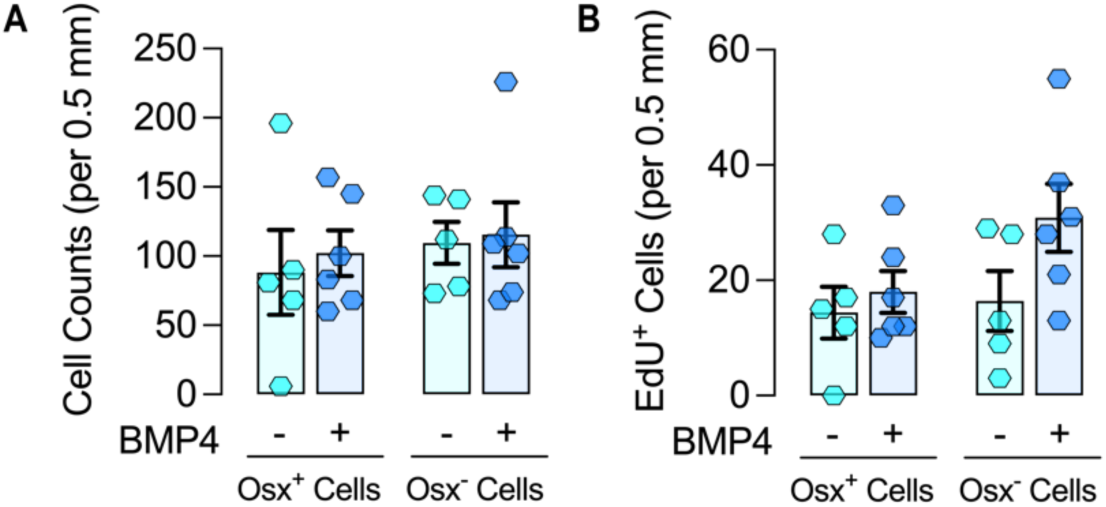
**A.** Total counts of Osx^+^ and Osx^−^ cells and **B.** proliferating Osx^+^ and Osx^−^ cells in the periosteum in YAP/TAZcKO^Osx^ mice treated with saline or BMP4. **E.** Total counts of proliferating Osx^+^ and Osx^−^ cells in the periosteum in WT^fl/fl^ and YAP/TAZcKO^Osx^ mice.

**Supplementary Figure 3:**
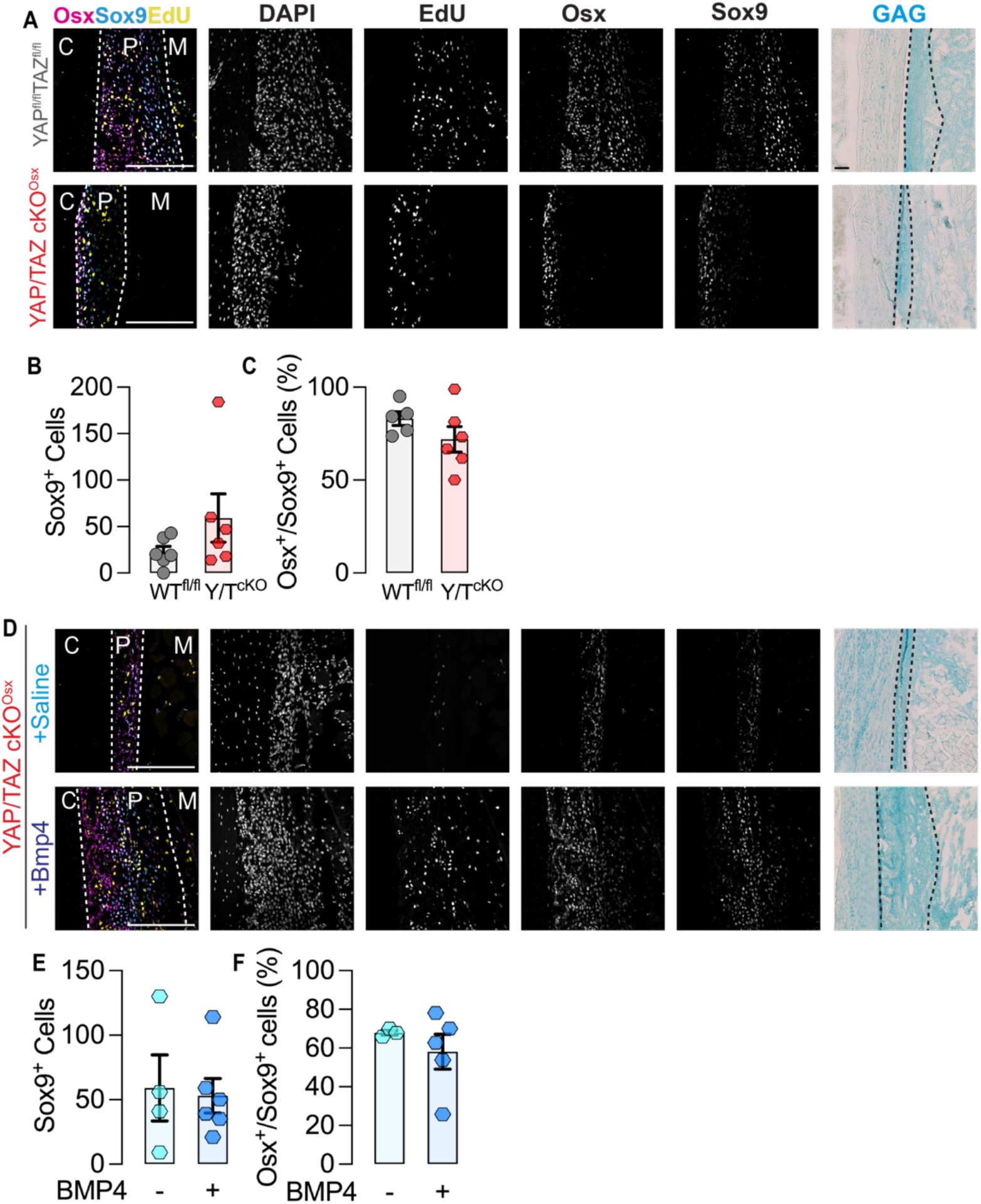
**A.** Staining for nuclei (DAPI), proliferating cells (EdU), immunostain for Osx and Sox9 and GAG staining in periostea of WT^fl/fl^ and YAP/TAZcKO^Osx^ mice. **B.** The number of Sox9+ cells per length of the periosteum was unchanged. **C.** Majority of these cells were positive for both Osx and Sox9. **D.** Staining for nuclei (DAPI), proliferating cells (EdU), immunostaining for Osx and Sox9 and GAG staining in periostea of YAP/TAZcKO^Osx^ mice treated with Saline or BMP4. **E.** The number of Sox9^+^ cells per length of the periosteum was unchanged. **F.** Majority of these cells were positive for both Osx and Sox9. Scale bar = 100 μM.

**Supplementary Figure 4:**
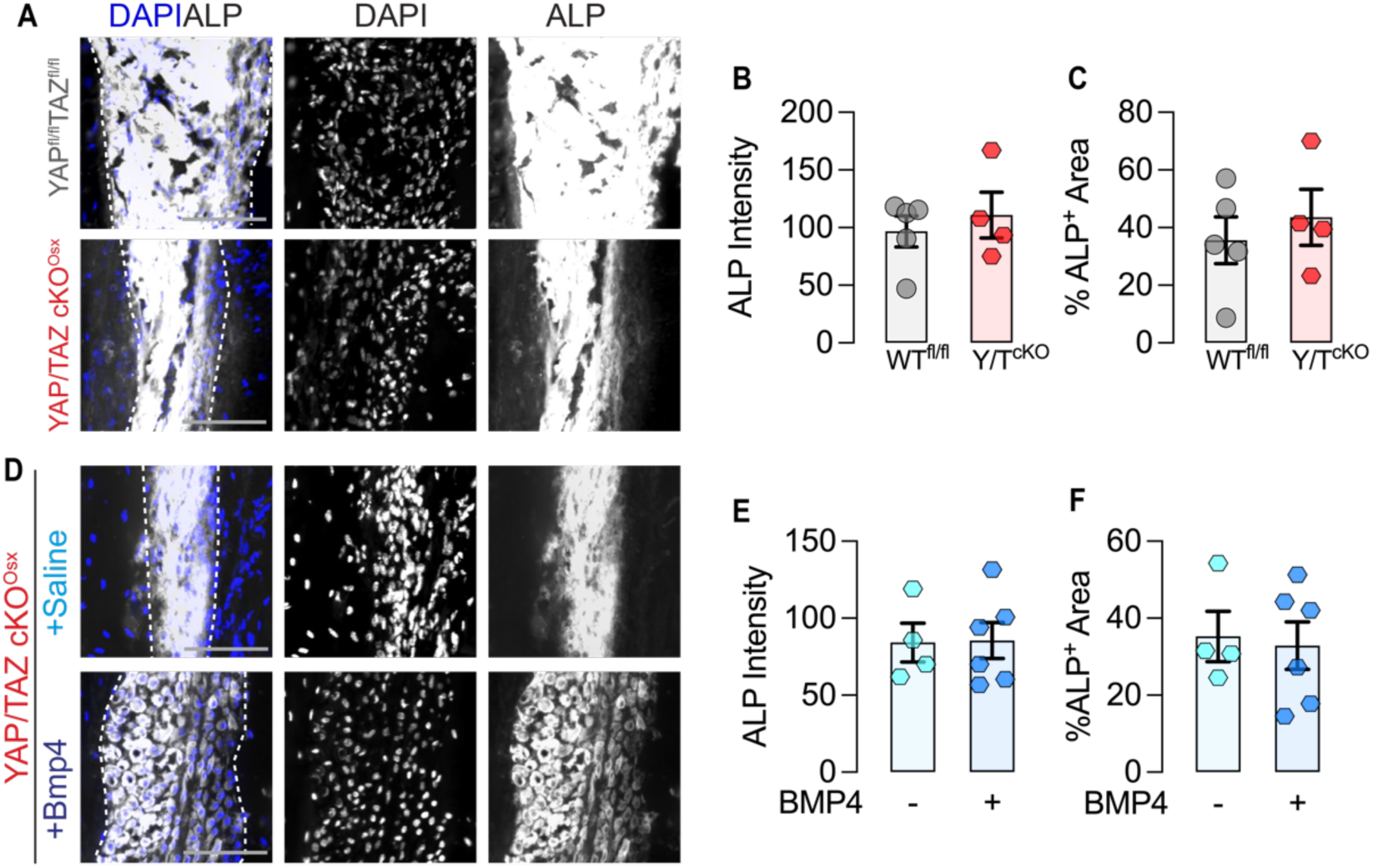
**A.** ALP activity and quantification of **B.** ALP intensity and **C.** % ALP^+^ area in the periosteum of WT^fl/fl^ and YAP/TAZcKO^Osx^ mice. **D.** ALP activity and quantification of **B.** ALP intensity and **C.** % ALP^+^ area in the periosteum of YAP/TAZcKO^Osx^ mice treated with saline and BMP4. Scale bar = 100 μM.

**Supplementary Figure 5:**
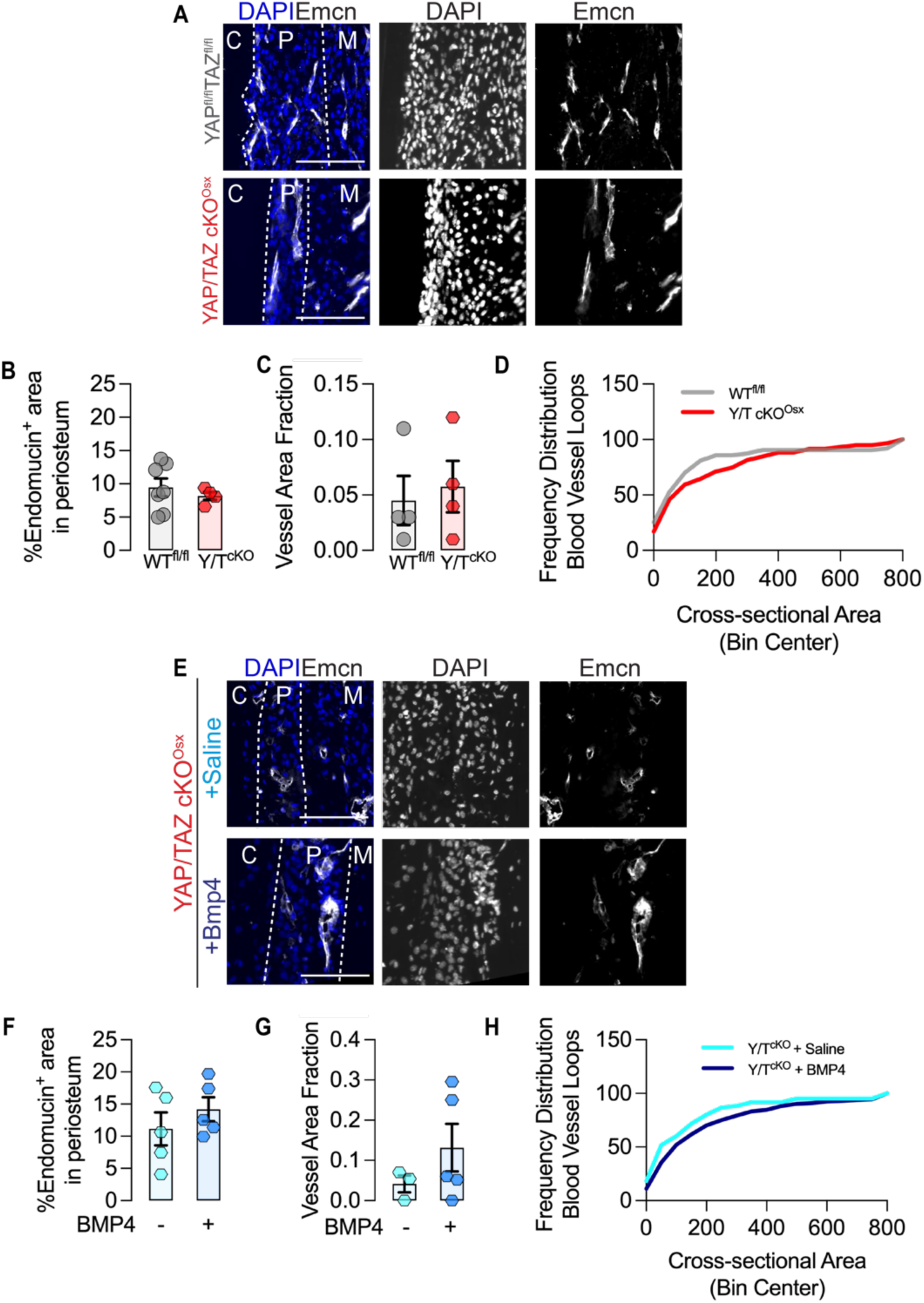
**A.** Endomucin staining for blood vessels, Quantification of **B.** Endomucin positive area, **C.** Vessel area fraction and **D**. Frequency distribution of blood vessel loops in the periosteum of WT^fl/fl^ and YAP/TAZcKO^Osx^ mice. **E.** Endomucin staining for blood vessels, Quantification of **F.** Endomucin positive area, **G.** Vessel area fraction and **H.** Frequency distribution of blood vessel loops in the periosteum of YAP/TAZcKO^Osx^ mice treated with saline or BMP4. Scale bar = 100 μM.

**Supplementary Figure 6:**
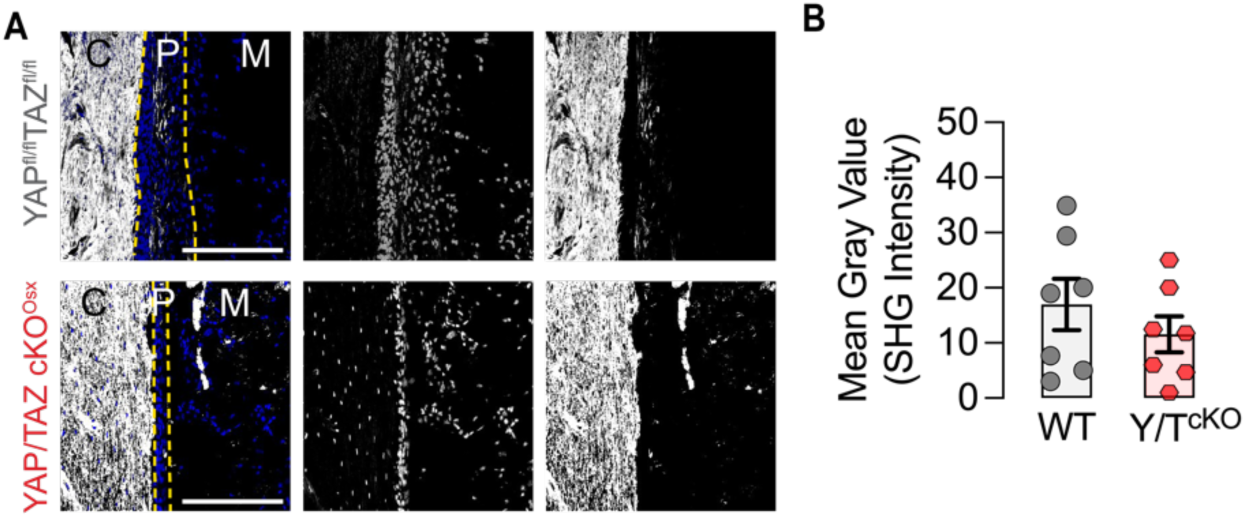
**A.** SHG imaging of periostea in WT^fl/fl^ and YAP/TAZcKO^Osx^ mice. **B.** Quantification of SHG intensity in the periostea of WT^fl/fl^ and YAP/TAZcKO^Osx^ mice. Scale bar = 100 μM.

## Notes

### Competing Interest Statement

The authors have declared no competing interest.

